# When Color Adds Nothing: A Causal Audit of Channel Triplication in Alzheimer’s MRI Classification

**DOI:** 10.64898/2026.07.07.737073

**Authors:** Singhvi Singhvi, Riddhi Singhvi

## Abstract

Medical imaging pipelines routinely copy single-channel grayscale data into three identical RGB channels before classification, usually without justification. This study tests whether that step affects model predictions. Four coordinated experiments on bit-identical RGB inputs sorted eleven classical machine learning models into three groups: five that were invariant to the copy, two that were nearly invariant, and four whose predictions changed.

On the Kaggle Alzheimer MRI Dataset (6,400 images, four classes, five seeds), five models (AdaBoost, HistGradientBoosting, KNN, SVM_Polynomial, and SVM_RBF) produced identical predictions in both conditions for every seed, where KNN is k-nearest neighbors and SVM a support vector machine, with polynomial and radial basis function (RBF) kernels. Two models (GaussianNB and SVM_Linear) differed by at most one of 1,280 samples, a dataset-dependent gap rather than exact invariance. The remaining four (DecisionTree, ExtraTrees, RandomForest, and LogisticRegression) differed substantively.

A regularization sweep on Logistic Regression traced its gap to a single cause. As L2 regularization weakened, the color-minus-grayscale macro F1 gap shrank steadily, from +12.07 percentage points at C=0.001 to near zero at C=100 (paired Wilcoxon p=0.0020 under strong regularization), showing the effect scales with feature count rather than image content.

A replication on the OASIS dataset, matched in size and class balance, reproduced every grouping, and the Logistic Regression gap reappeared in the same direction at smaller magnitude (+5.30 points macro F1). Two deep networks, ResNet18 and DenseNet121, gave identical predictions across all twenty paired conditions. Channel triplication left most models unchanged while multiplying classical training time 2.3 to 4.0 times without benefit.

## 1. Introduction

In 2024, an estimated 6.9 million Americans aged 65 or older were living with Alzheimer’s disease (Alzheimer’s Association, 2024). The condition is the seventh leading cause of death in the United States and one of the most costly. Early detection allows earlier intervention, which is associated with better clinical outcomes. Magnetic resonance imaging (MRI) is among the most accessible non-invasive diagnostic tools, and a growing body of research applies machine learning to MRI-based classification of Alzheimer’s disease (Kumar & Shastri, 2022; Liu et al., 2014; Khan et al., 2019). The Kaggle Alzheimer MRI Image Dataset has emerged as a common benchmark for this work.

A standard preprocessing step in this literature is channel triplication: a single-channel grayscale MRI is converted to a three-channel RGB image by copying grayscale values across R, G, and B. The common implicit assumption is that triplication is a no-op: the model either ignores the redundancy or correctly identifies it. This assumption has not been systematically tested. Prior work on input redundancy in medical imaging has focused on data augmentation strategies (Shorten & Khoshgoftaar, 2019) and on pretrained model fine-tuning (Raghu et al., 2019), without isolating the channel triplication step itself. Reproducibility audits of medical imaging classifiers have noted variability across seeds and preprocessing choices (McDermott et al., 2021), without identifying channel triplication as a specific source of artifact.

This research presents a four-phase empirical investigation. Phase 1 trained 11 classical machine learning models under grayscale and color input conditions across five random seeds on the Kaggle dataset (110 cells) and partitioned the models into empirical groups based on byte-level prediction comparisons. Phase 2 tested the proposed Logistic Regression mechanism through a controlled regularization ablation, running a six-point regularization sweep across ten seeds per condition (120 cells) on the same dataset. Phase 3 tested external validity by replicating Phase 1 on the OASIS Alzheimer’s Detection dataset (110 cells), subsampled to match the Phase 1 scale and class distribution exactly. Phase 4 tested whether the partition extends to deep learning by training ResNet18 and DenseNet121 across both datasets, both channel conditions, and five seeds (40 cells).

The four experiments together show that, for the classifiers and datasets tested, channel triplication has classifier-dependent effects mediated by specific mathematical properties of each model’s training procedure rather than by the data itself. Five of the 11 classical models are structurally invariant to triplication. Four of the 11 are systematically affected by two distinct mechanisms. The remaining two of the 11 are empirically near-invariant: they differ by at most one sample on Kaggle and not at all on OASIS, a dataset-dependent near-invariance that is not guaranteed to be exact (Section 5). Two findings support the Logistic Regression result. First, a controlled regularization ablation showed the color-minus-grayscale gap shrinking steadily as regularization weakened, pointing to regularization strength as the cause rather than any property of the data. Second, the full pattern held on a second dataset, indicating that the results are not specific to a single dataset.

## 2. Objectives and Hypotheses

This research tests five hypotheses:

**H1.** Classical machine learning models partition into invariance groups under channel triplication on bit-identical RGB inputs. The partition is determined by the mathematical structure of each model’s training procedure and predictions.

**H2.** Some models base their decisions solely on pixel values, distances between data points, or feature rankings. For these models, the input being grayscale or color makes no difference; they produce identical predictions in both cases.

**H3.** In Logistic Regression, the gap between color and grayscale performance, measured by accuracy and macro F1, depends on the strength of the L2 regularization. As the parameter C grows larger and the regularization weakens, the gap steadily shrinks, approaching zero once the regularization term contributes negligibly to the outcome.

**H4.** The invariance partition observed in one dataset replicates in another independent dataset with matched scale and class distributions. Direction and rank order of effects are preserved across datasets; effect magnitudes depend on the data distribution.

**H5.** The group of perfectly identical models also includes deep convolutional networks (CNNs) trained on three-channel input. When a grayscale image is copied into three identical channels to match the network’s expected input shape, the result is numerically identical to an image loaded as color from the start. Because the input is identical, the trained network produces identical predictions either way.

## 3. Materials and Methods

### 3.1 Datasets

This study uses two datasets. The first is the Kaggle Alzheimer MRI Image Dataset (Kumar & Shastri, 2022), which contains 6,400 brain MRI images across four diagnostic classes: Non Demented (n=3,200; 50.0%), Very Mild Demented (n=2,240; 35.0%), Mild Demented (n=896; 14.0%), and Moderate Demented (n=64; 1.0%).

The second is the OASIS Alzheimer’s Detection dataset (Aithal, 2024), derived from the Open Access Series of Imaging Studies (Marcus et al., 2007) and based on 461 patients with Clinical Dementia Rating annotations. It contains approximately 86,400 brain MRI slices across the same four classes.

For the Phase 3 cross-dataset replication, the OASIS dataset was deterministically subsampled (seed=0) to 6,400 images matching the Kaggle dataset’s class distribution exactly (3,200 / 2,240 / 896 / 64). This subsampling isolated dataset effects (patient cohort, acquisition protocol, slice selection) while holding sample size and class distribution constant relative to the Kaggle Phase 1 experiment, so that any observed difference between the two datasets is attributable to the data itself rather than to confounding factors of scale or class balance. Because the subsample is deterministic, it can be reproduced exactly, and the full manifest is reported in the data availability section.

### 3.2 Channel Identity Verification

Before any classification experiments on each dataset, the channel content of every image was verified at the pixel level using integer arithmetic. For each image, the maximum absolute difference between the R-G, R-B, and G-B channels was computed. An image was considered to have distinct channels if any of these differences was greater than zero.

The result was the same for both datasets. On the Kaggle dataset, all 6,400 images had bit-identical channels: no image had distinct channels, and the maximum R-G, R-B, and G-B differences were all zero. The OASIS subsample yielded the same outcome, with all 6,400 images bit-identical and every maximum difference equal to zero.

Although both datasets are stored in three-channel RGB format, every image therefore carries no more information than a single grayscale channel.

### 3.3 Experimental Design

#### Preprocessing

All images were resized to 128 × 128 pixels using bilinear interpolation and then flattened. For grayscale input, images were converted to single-channel mode before resizing, yielding 16,384 features per image. For color input, the three-channel mode was retained, yielding 49,152 features per image. Within each image, the color features are simply three identical copies of the grayscale features. Each dataset was divided into training (60%), validation (20%), and test (20%) sets using stratified shuffle splits across multiple random seeds, and features were normalized to the unit interval using a MinMaxScaler fitted to the training set.

#### Experimental design

The study ran in four phases. Phase 1 evaluated 11 classifier families (AdaBoost, DecisionTree, ExtraTrees, GaussianNB, HistGradientBoosting, KNN, LogisticRegression, RandomForest, SVM_Linear, SVM_Polynomial, and SVM_RBF), where KNN denotes k-nearest neighbors and SVM_Linear, SVM_Polynomial, and SVM_RBF denote support vector machines (SVMs) with linear, polynomial, and radial basis function (RBF) kernels, on the Kaggle dataset across five seeds and both channel conditions, producing a 110-cell matrix. Phase 2 fixed the classifier to Logistic Regression and varied the regularization parameter C over six values ({0.001, 0.01, 0.1, 1, 10, 100}), again across ten seeds and both channel conditions, on the Kaggle dataset, for a 120-cell matrix. Phase 3 repeated the Phase 1 design (11 classifiers, two channels, five seeds) on the OASIS subsample, giving a second 110-cell matrix. Phase 4 evaluated two ImageNet-pretrained convolutional architectures, ResNet18 and DenseNet121, across five seeds and both channel conditions on both datasets, producing a 40-cell matrix.

#### Deep learning setup

In Phase 4, grayscale input was loaded as a single channel and expanded to three identical channels by tensor repetition, while color input was loaded as native three-channel RGB. Each network was fine-tuned for 12 epochs using Adam (learning rate 1e-3, weight decay 1e-4) with a batch size of 64. Determinism was enforced throughout, with seeded initialization, deterministic cuDNN, and seeded DataLoader ordering.

#### Software

Phases 1 through 3 used scikit-learn 1.3.2; Phase 4 used PyTorch 2.1 with torchvision. All experiments were run under Python 3.10.9. Hyperparameters are summarized in Table 4.

### 3.4 Evaluation Metrics

Each trained model was evaluated on its corresponding test set using test accuracy, balanced accuracy, macro F1, Cohen’s kappa, and per-class F1 and recall. For test accuracy, 95% confidence intervals were estimated using nonparametric bootstrap resampling with 500 replications. For the channel-invariance analysis, predictions on the same test set under grayscale and color conditions were compared at the per-sample level, recording the number of samples where the two predictions differed.

### 3.5 Statistical Analysis

For each model and metric, paired bootstrap resampling with 10,000 replications on the seed-paired differences produced a 95% confidence interval on the color-minus-grayscale mean difference. Paired Wilcoxon signed-rank tests were computed on the same data. The Wilcoxon test has a floor on the size of its p-value, set by the number of paired observations. With five pairs, the smallest achievable two-sided p-value is 2/2⁵ = 0.0625, reached when all differences share the same sign. The Phase 2 sweep used ten pairs per condition, which lowers this floor to 2/2¹⁰ = 0.00195 ≈ 0.0020. Bootstrap confidence intervals are reported as the primary statistical evidence, and Wilcoxon p-values are reported as supplementary.

## 4. Results

### 4.1 Channel Identity Confirmation

All 6,400 images in the Kaggle dataset and the OASIS subsample contain bit-identical R, G, and B channels. No images in either dataset had any pixel for which channel values differed. The grayscale and color inputs therefore carry the same information at the pixel level; the only difference is the feature count (16,384 for grayscale versus 49,152 for color).

### 4.2 Phase 1: Three Empirical Groups in the Kaggle Dataset

Across the 110 Phase 1 cells, the 11 models fell into three empirical groups, based on a byte-level comparison of their predictions.

Five models produced byte-identical predictions across both channel conditions for all five seeds: AdaBoost, HistGradientBoosting, KNN, SVM_Polynomial, and SVM_RBF. On the 1,280-sample test set, their prediction vectors matched at every position, and all metrics (test accuracy, macro F1, balanced accuracy, Cohen’s kappa, and per-class scores) agreed to machine precision.

Two models were near-identical. GaussianNB differed by a single sample in two of five seeds, and SVM_Linear differed by a single sample in one of five seeds. In each case, the lone disagreement fell at a class-margin boundary. As Section 5 explains, these are dataset-dependent near-misses rather than evidence of exact structural invariance.

Four models produced substantively different predictions: DecisionTree (171 to 215 disagreements per seed), RandomForest (106 to 144), ExtraTrees (67 to 85), and LogisticRegression (50 to 72). Two distinct mechanisms drive these differences within the divergent group, as explained below.

Figure 1 shows test accuracy across the 11 models. Figure 2 shows the per-seed byte-level disagreement counts.

**Figure 1.**
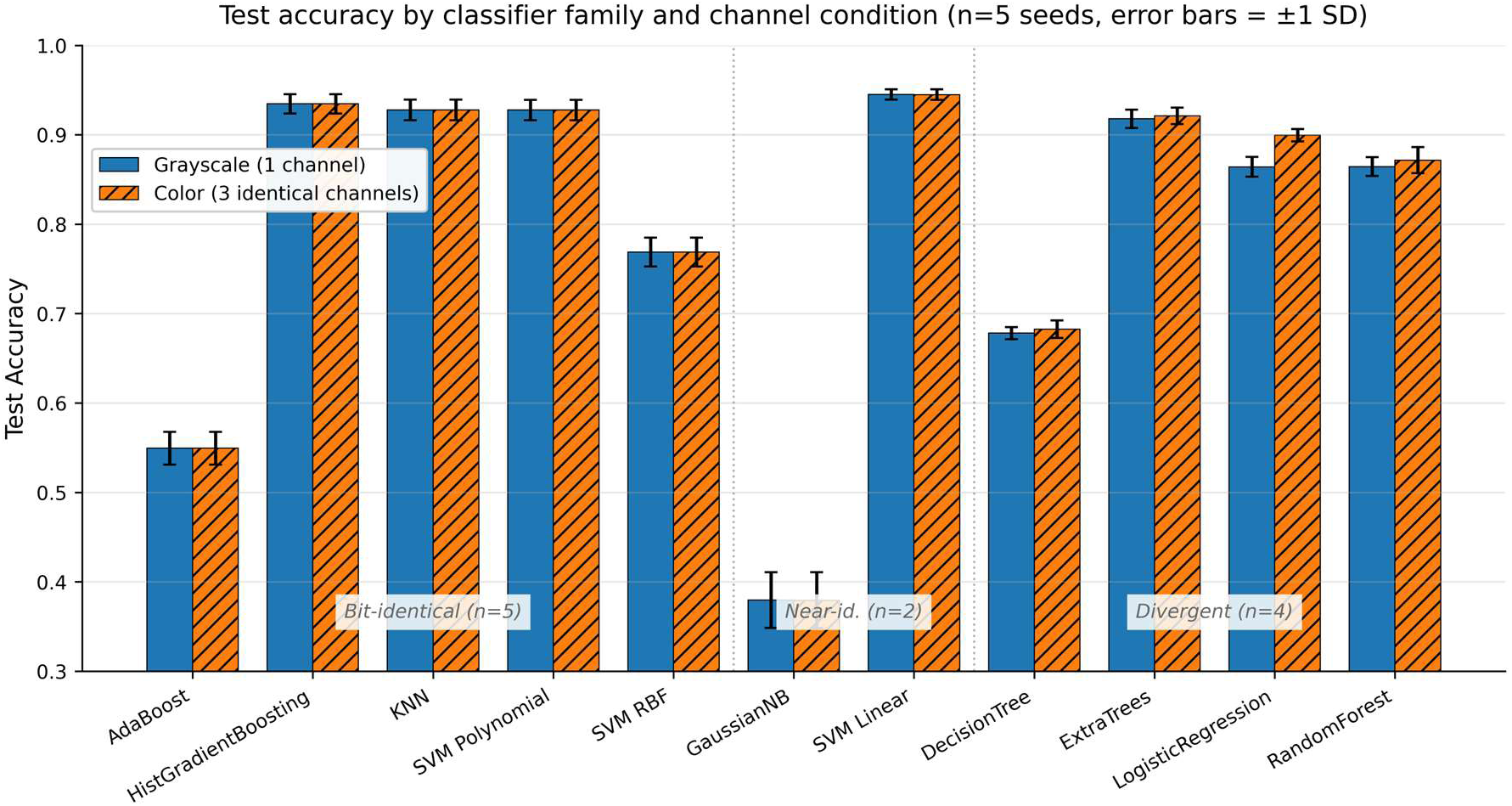
Phase 1 test accuracy by classifier family and channel condition (Kaggle dataset, n=5 seeds, error bars = ±1 SD). The 11 models partition into three empirical groups: bit-identical (n=5, left), near-identical (n=2, center), and divergent (n=4, right). Within bit-identical and near-identical groups, color and grayscale bars are visually indistinguishable; within the divergent group, color differs from grayscale by a small to moderate margin depending on the underlying mechanism.

**Figure 2.**
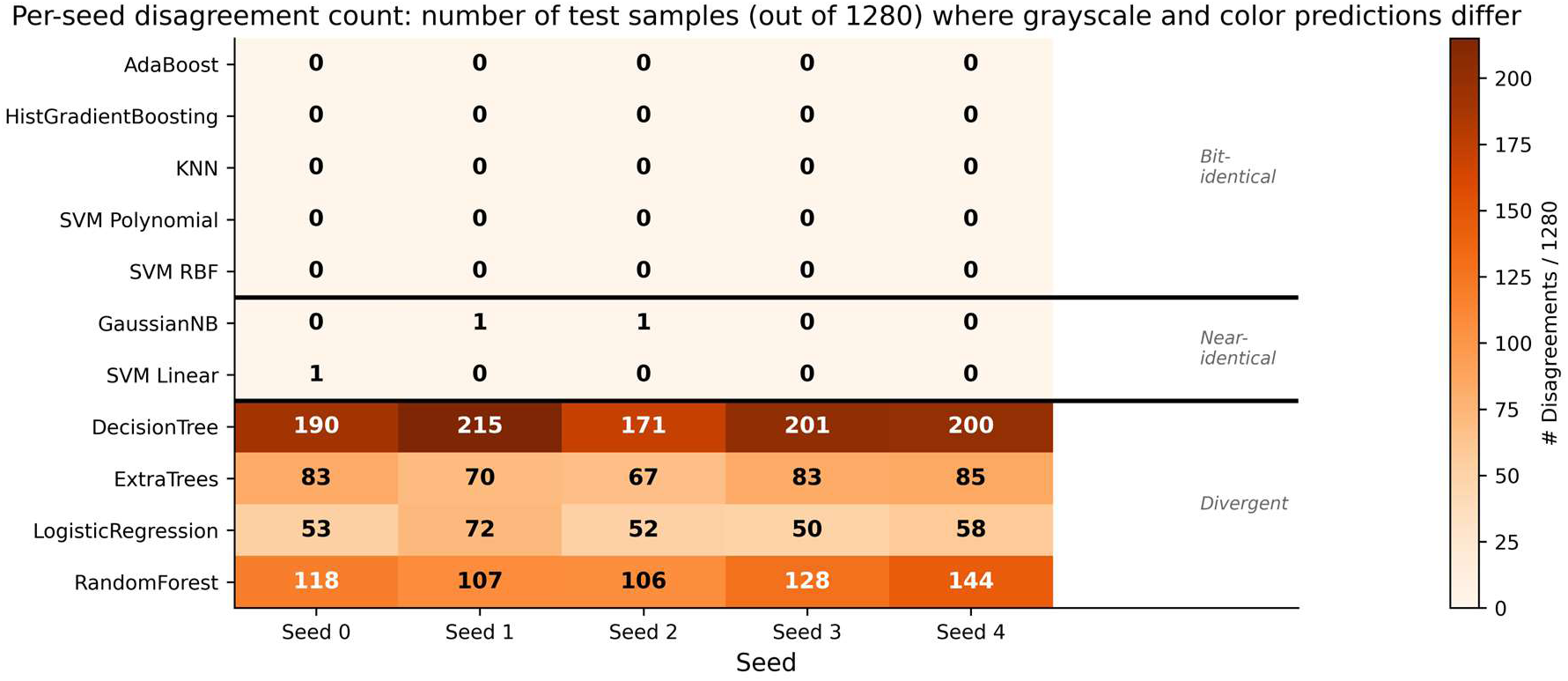
Phase 1 per-seed prediction disagreement count on the Kaggle dataset (number of test samples out of 1,280 where grayscale and color predictions differ). Bit-identical models show zero disagreements at all seeds; near-identical models show at most one disagreement; divergent models show 50 to 215 disagreements per seed depending on the underlying mechanism.

### 4.3 Two Mechanisms Within the Divergent Group

The four divergent models do not all diverge for the same reason. Two distinct mechanisms are at work.

The first is a coupling between the random-number generator (RNG) and the number of features. It affects DecisionTree, RandomForest, and ExtraTrees through two related routes. RandomForest and ExtraTrees draw a random subset of candidate features at each split (with max_features=’sqrt’, that is, 128 candidates from the 16,384 grayscale features, versus 221 from the 49,152 color features). Even with the random state fixed, a seeded draw from a 49,152-element index space selects a different subset than the equivalent draw from a 16,384-element space, so different splits, trees, and predictions result. DecisionTree, at scikit-learn defaults (max_features=None), does not subsample; it evaluates every feature at each split, so no information is lost. Its divergence comes instead from the RNG-shuffled order in which the splitter visits features. That order is consumed differently at 49,152 versus 16,384 features, and it governs which split is kept when several candidates yield equal impurity improvement, so tie resolution, and therefore the resulting tree, changes with the feature count.

The key point is this: in both routes the information available at each split is identical, since the extra features are exact copies of the originals. What differs is the model’s stochastic path through that information. This is the stochastic mechanism.

The effects are small and non-systematic. Across the five Phase 1 seeds, color input gave slightly higher accuracy than grayscale: 0.72 points for RandomForest, 0.44 for DecisionTree, and 0.33 for ExtraTrees. In every case, the paired bootstrap 95% confidence interval spans zero.

The second mechanism, Tikhonov coupling, affects LogisticRegression. Its training objective with L2 regularization minimizes the negative log-likelihood plus a penalty equal to the squared L2 norm of the weight vector, divided by twice the regularization parameter C. When features are triplicated, the optimizer can spread the same target weight across three identical copies, giving each copy one-third of the magnitude. This redistribution reduces the squared norm of the weight vector, effectively relaxing the per-feature regularization.

At C=0.01 in Phase 1, this relaxation produced a substantively different optimum. Compared with grayscale, color input improved accuracy by 3.55 points, macro F1 by 7.44 points, and minority-class recall by 26.2 points, with paired bootstrap 95% confidence intervals excluding zero in all three cases. The minority-class figure should be read with caution, given the very small minority-class test sample (Moderate Demented, n=13 per seed); Section 6.4 discusses this limitation in detail. Phase 2 tests this mechanism directly through a controlled regularization ablation.

### 4.4 Phase 2: Regularization Sweep Supports the Tikhonov Mechanism

Phase 2 varied the Logistic Regression regularization parameter C across six values, from 0.001 to 100, holding everything else constant. The Tikhonov mechanism makes a clear prediction: as C increases and regularization weakens, the color-minus-grayscale gap should shrink steadily and approach zero once the L2 term barely affects the objective. Figure 3 shows the results for the three headline metrics. The observed pattern closely matches this prediction. The gap shrank monotonically toward zero as C increased, providing strong evidence for the Tikhonov mechanism from a controlled ablation.

**Figure 3.**
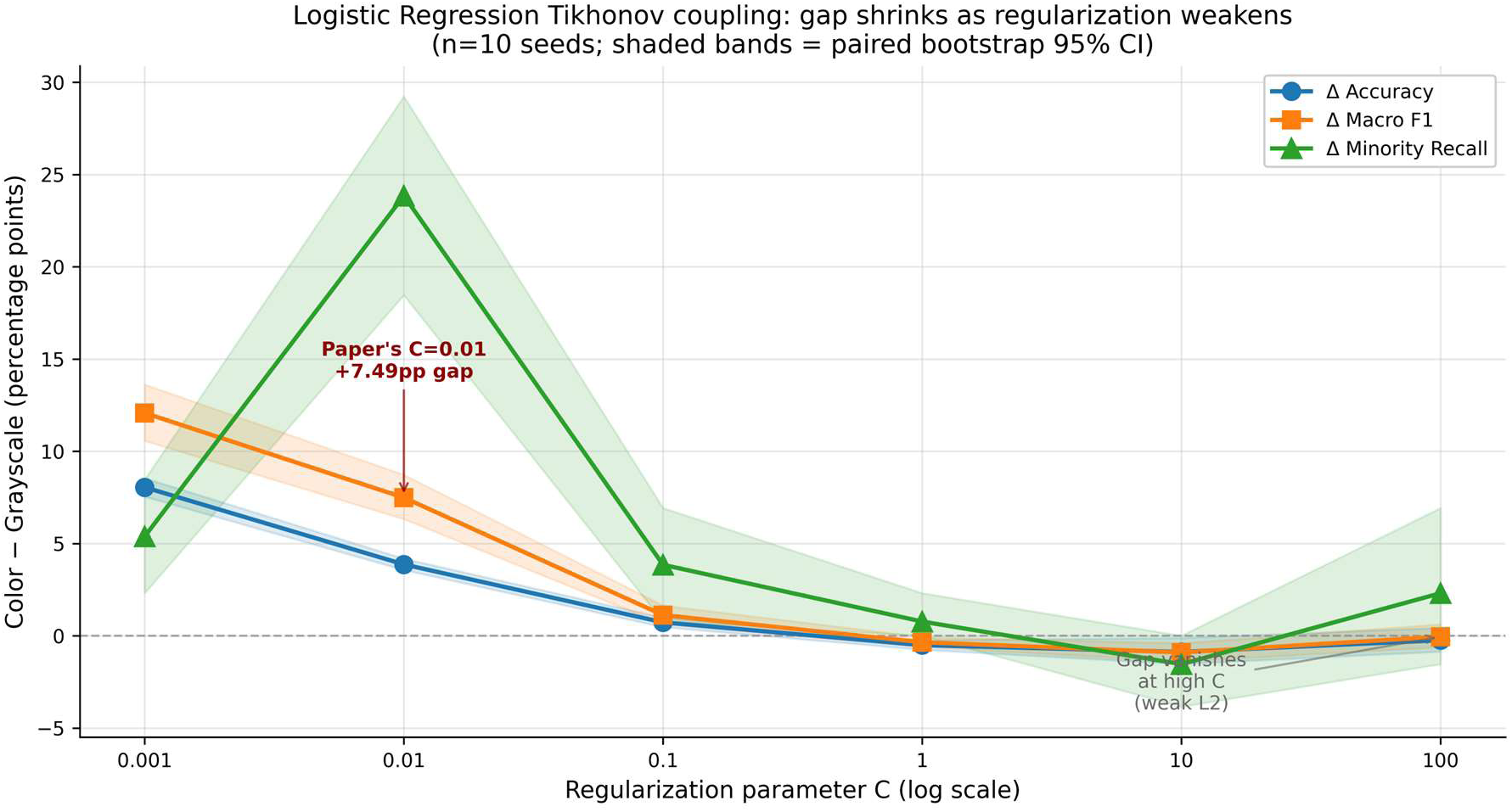
Phase 2 Logistic Regression color-minus-grayscale gap across six values of the regularization parameter C (Kaggle dataset, n=10 seeds per condition, shaded bands = paired bootstrap 95% CI). Under strong regularization (C=0.001 to C=0.1), color produces systematically higher accuracy, macro F1, and minority-class recall than grayscale, with confidence intervals excluding zero. As regularization weakens (C ≥ 1), the gap shrinks toward zero. At C=100, the gap is -0.05 percentage points on macro F1 with a confidence interval that spans zero. The pattern is consistent with the Tikhonov coupling prediction: the color-versus-grayscale effect in Logistic Regression is mediated by L2 regularization scaling with feature count.

The macro F1 gap shrank from +12.07 percentage points at C=0.001 (paired bootstrap 95% CI [+10.56, +13.61]) to +7.49 points at C=0.01 (CI [+6.32, +8.72]), to +1.12 points at C=0.1 (CI [+0.63, +1.64]), and to within 0.05 points of zero at C=100 (CI [-0.62, +0.62]). The same monotonic shrinkage held for accuracy and minority-class recall. With ten paired seeds, paired Wilcoxon signed-rank tests yielded p-values of 0.0020 at the strong-regularization conditions, below the conventional α=0.05 threshold. The Phase 2 estimates at C=0.01 use ten seeds, whereas the Phase 1 and Table 3 estimates at the same C use five; the two agree within seed-sampling noise (+7.49 vs +7.44 pp macro F1, +23.85 vs +26.2 pp minority-class recall, +3.87 vs +3.55 pp accuracy).

A subtle secondary finding emerged between C=1 and C=10. In this range of weak regularization, color produced slightly worse accuracy and macro F1 than grayscale (Δ macro F1 from -0.35 to -0.91 points), with confidence intervals narrowly excluding zero. The likely cause is numerical: with light L2 regularization, the optimization landscape is dominated by the data-fit term, and triplicated features raise the dimensionality without adding information, slightly degrading the solver’s conditioning. At very weak regularization (C=100), this conditioning effect itself vanishes, and the two conditions converge.

Figure 4 shows the minority-class recall trajectory across the same six C values, separated by condition. Under strong regularization, color substantially outperforms grayscale on minority-class recall, with high seed-to-seed variance in the grayscale condition. As C increases, the two conditions converge.

**Figure 4.**
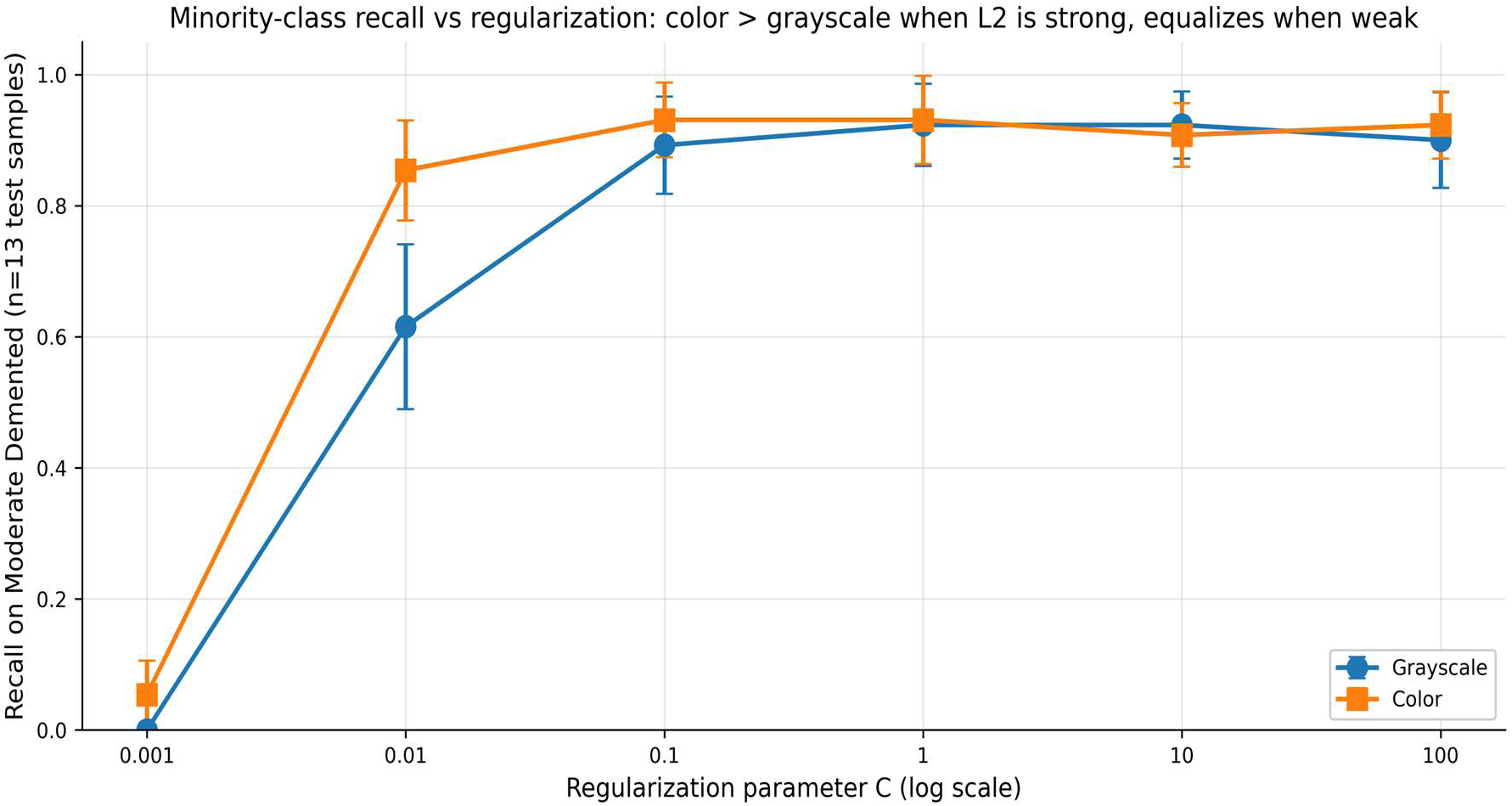
Phase 2 minority-class (Moderate Demented, n=13 test samples) recall by C and channel condition (n=10 seeds, error bars = ±1 SD). Under strong regularization (C=0.001 to C=0.01), color produces substantially higher minority-class recall than grayscale, with the gap reaching 23.85 percentage points at C=0.01. As regularization weakens (C ≥ 0.1), the two conditions converge.

### 4.5 Phase 3: Cross-Dataset Replication on OASIS

Phase 3 replicated the Phase 1 design on the OASIS Alzheimer’s Detection dataset, subsampling to match the Phase 1 scale and class distributions exactly. The OASIS subsample passed the same channel-identity verification: zero images with distinct R, G, or B channels across all 6,400 subsampled images, with all maximum pairwise channel differences equal to zero. The 11 models were trained with five seeds per condition using both grayscale and color inputs. Per-seed prediction disagreement counts for both datasets are shown in Figure 5.

**Figure 5.**
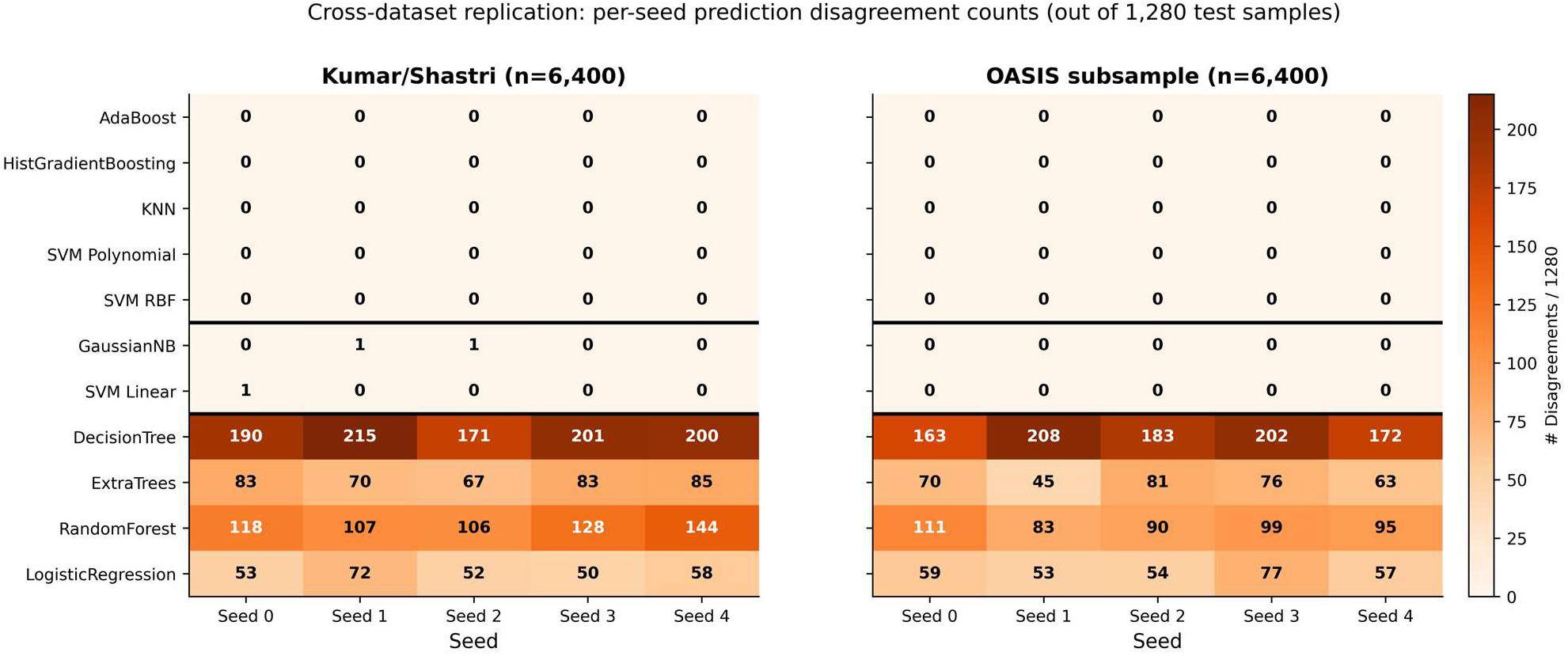
Cross-dataset replication: per-seed prediction disagreement counts (out of 1,280 test samples) on the Kaggle dataset (left) and the OASIS subsample (right). The five Kaggle-bit-identical models remain bit-identical on OASIS. The four Kaggle-divergent models remain divergent on OASIS with similar per-seed magnitudes. The two Kaggle-near-identical models (GaussianNB, SVM_Linear) produced zero disagreements on OASIS, indicating that their near-invariance is empirical and dataset-dependent rather than a proven structural property.

All five Kaggle-bit-identical models remained invariant on OASIS: AdaBoost, HistGradientBoosting, KNN, SVM_Polynomial, and SVM_RBF produced zero prediction disagreements across all seeds. All four Kaggle-divergent models remained divergent: DecisionTree (163 to 208 disagreements per seed), RandomForest (83 to 111), ExtraTrees (45 to 81), and LogisticRegression (53 to 77). The per-seed disagreement magnitudes were similar to those on Kaggle, consistent with the stochastic and Tikhonov mechanisms being properties of the algorithm and feature space rather than of the data distribution.

The two Kaggle-near-identical models, GaussianNB and SVM_Linear, produced zero disagreements on OASIS for all five seeds. Their near-invariance is therefore empirical and dataset-dependent rather than a proven structural property. The full partition is now clear: five classifiers (AdaBoost, HistGradientBoosting, KNN, SVM_Polynomial, SVM_RBF) are structurally invariant under channel triplication, four (DecisionTree, ExtraTrees, RandomForest, LogisticRegression) are divergent, and two (GaussianNB, SVM_Linear) are empirically near-invariant for the reasons given in Section 5. Treated as a practical no-op group, the five structurally invariant models together with the two near-invariant models produced at most one-sample disagreements on Kaggle and none on OASIS.

The Logistic Regression Tikhonov effect replicated on OASIS in the same direction but with reduced magnitude. Color accuracy exceeded grayscale by 2.52 percentage points (versus 3.55 on Kaggle), macro F1 by 5.30 (versus 7.44), and minority-class recall by 10.77 (versus 26.2). The reduced magnitude is consistent with the mechanism: the effect’s direction is structural, set by L2 coupling to the feature count. At the same time, its size depends on the extent to which L2 regularization influences classification on a given dataset. Three of the four divergent models show the same direction of effect on both datasets for the reported metrics; the fourth (DecisionTree per-class results) shows non-systematic variation, as expected from the stochastic mechanism.

### 4.6 Computational Cost

Channel triplication increased wall-clock training time by 2.3 to 4.0 times for the same classical model across all three classical-ML phases, depending on the algorithm. The increase reflects the threefold rise in feature count. For Logistic Regression at C=0.01 on the Kaggle dataset, color training took 1,987 seconds per seed, compared with 509 seconds for grayscale, a 3.9-fold increase. For HistGradientBoosting, color required 774 seconds, compared with 266 for grayscale, a 2.9-fold increase. Across the five bit-identical classical models on Kaggle, this extra compute was pure overhead: the color condition produced byte-identical predictions to grayscale at every seed. The full 110-cell Phase 1 experiment took 9.7 hours of wall-clock time on a single CPU instance, whereas the grayscale-only subset would have taken approximately 2.0 hours.

Convolutional networks tell a different story. In Phase 4, both the grayscale and color code paths produce a (3, H, W) input tensor, where 3 is the number of channels and H and W are the image height and width in pixels, so training times are nearly identical (ResNet18 on Kaggle: 1.09 minutes grayscale versus 1.08 minutes color; DenseNet121 on OASIS: 2.91 minutes grayscale versus 2.97 minutes color). The classical-ML compute waste therefore does not apply to the standard deep-learning setup.

### 4.7 Phase 4: Deep Convolutional Architectures

Phase 4 tested whether the partition observed in classical machine learning also holds in deep learning. ResNet18 and DenseNet121, both ImageNet-pretrained, were trained on both datasets under both channel conditions, with five seeds per cell, for a total of 40 training runs. The grayscale code path loaded each image as a single channel and then expanded it to three identical channels by tensor repetition before feeding the network. The color code path loaded each image as native three-channel RGB. For images with bit-identical R, G, and B channels, the two paths produce numerically identical (3, H, W) input tensors. Each network was trained for 12 epochs with Adam (learning rate 1e-3, weight decay 1e-4), batch size 64, at 128 × 128 resolution, with deterministic seeded initialization, deterministic cuDNN, and seeded DataLoader ordering enforced throughout.

In all 20 paired (architecture, dataset, seed) conditions, grayscale and color predictions agreed byte-for-byte, and the per-seed disagreement count was zero in every cell. Test accuracy, macro F1, balanced accuracy, Cohen’s kappa, and every per-class metric matched between conditions to machine precision. ResNet18 reached a mean test accuracy of 0.9358 on Kaggle and 0.9244 on the OASIS subsample, identical across channel conditions; DenseNet121 reached 0.9244 on Kaggle and 0.9134 on OASIS, again identical across conditions. Per-seed accuracies showed normal seed-to-seed variation, but the grayscale and color seed-paired values were identical at every position (Figure 6).

**Figure 6.**
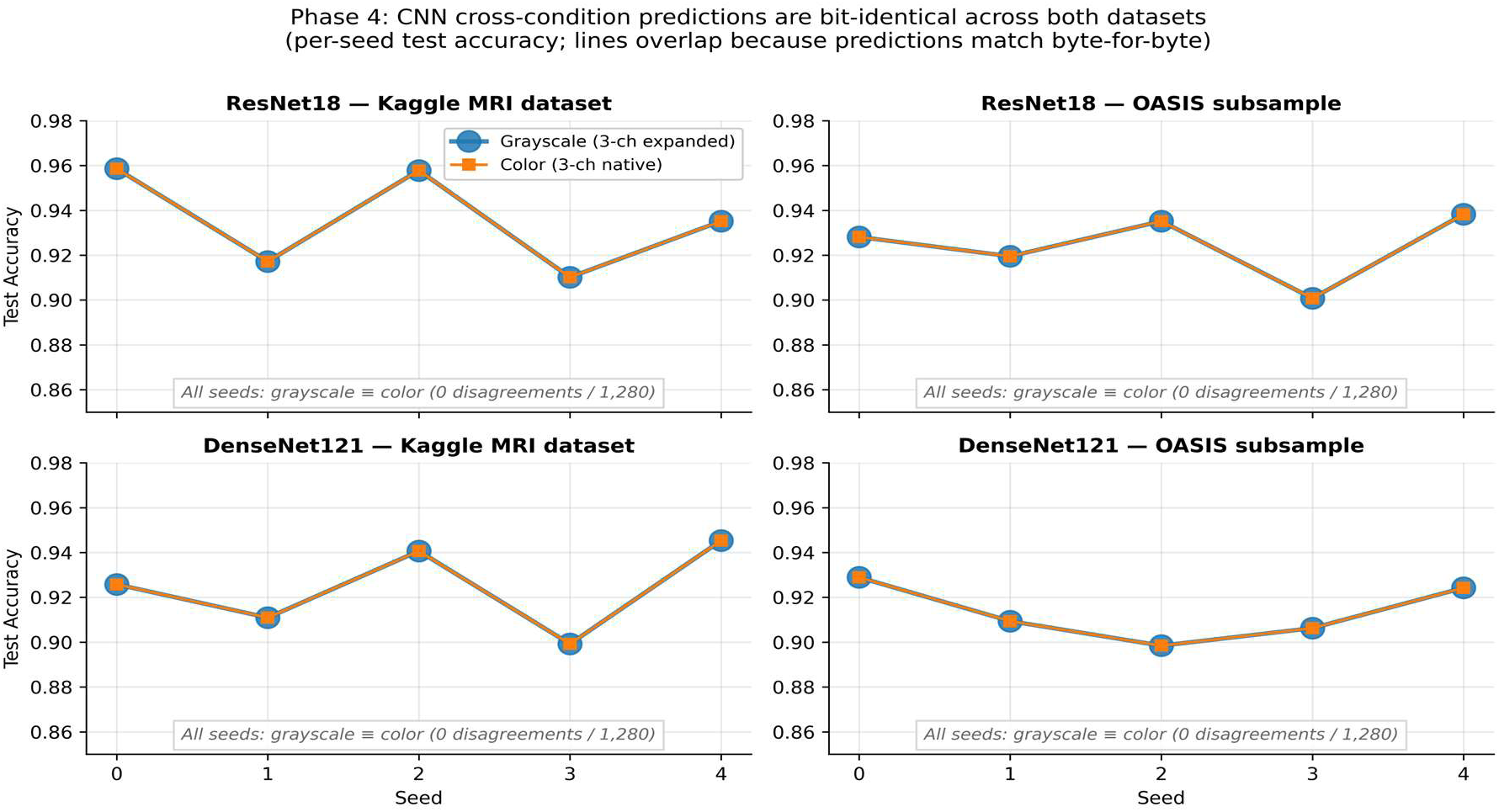
Phase 4 CNN test accuracy by architecture, dataset, and channel condition (n=5 seeds per cell). Grayscale (blue circles) and color (orange squares) markers sit at the same position for every seed in all four panels, reflecting byte-identical predictions in all 20 paired cells. The bit-identical group of the framework extends from classical machine learning to ImageNet-pretrained convolutional networks with channel-triplicated input.

This result extends the bit-identical group beyond the seven classical models to include deep convolutional networks under the standard transfer-learning setup. The reason is straightforward. When grayscale data is expanded to three identical channels to match a network’s expected input shape, the resulting input tensor is numerically identical to a natively loaded triplicated-RGB tensor. Given identical inputs, identical initialization seeds, identical training-data ordering, and deterministic cuDNN operations, the trained networks and their predictions come out byte-identical. This invariance is expected for architectures trained under the same deterministic setup. Other configurations, such as custom single-channel input convolutions, vision transformers, or non-pretrained architectures, were not tested, so the result covers the dominant deep-learning paradigm in medical imaging rather than every possible setting. An ImageNet-pretrained convolutional network fine-tuned on grayscale data and triplicated to RGB is, at the prediction level, invariant to whether the triplication is performed via tensor expansion or native RGB loading.

An earlier exploratory expectation was that these models might fall into the divergent group because CNNs could diverge due to stochastic filter initialization. That expectation was wrong. The stochastic mechanism applies only when an algorithm’s internal seeded subsampling depends on the feature count, as is the case for the four classical divergent models. For CNNs in the standard transfer-learning setup, the feature count and input tensor shape are identical between conditions, so the mechanism does not apply. This failed expectation is reported openly because the corrected understanding is the more useful one for the field: the standard practice of triplicating grayscale medical imaging data to fit pretrained color CNNs is, at the prediction level, a no-op.

### 4.8 Summary Tables

Table 1 reports the Phase 2 regularization sweep results in compact form. Table 2 reports the cross-dataset invariance classification. Table 3 reports the Logistic Regression effect sizes on both datasets.

**Table 1.** Phase 2 Logistic Regression color-minus-grayscale gap by C value (Kaggle dataset, n=10 seeds per condition). Δ values are mean differences in percentage points; 95% CIs are paired bootstrap intervals. Under strong regularization (C=0.001 to C=0.1), the gap is large, and CIs exclude zero. At weak regularization (C ≥ 1), the gap is small with CIs that span zero on minority recall.

| C | $\Delta$ Acc (pp) | $\Delta$ Acc 95% CI | $\Delta$ Macro F1 (pp) | $\Delta$ F1 95% CI | $\Delta$ Min Rec (pp) | $\Delta$ Min Rec 95% CI |
| --- | --- | --- | --- | --- | --- | --- |
| 0.001 | +8.05 | [+6.18, +9.97] | +12.07 | [+10.56, +13.61] | +5.39 | [-2.69, +13.85] |
| 0.01 | +3.87 | [+3.39, +4.42] | +7.49 | [+6.32, +8.72] | +23.85 | [+17.69, +30.00] |
| 0.1 | +0.73 | [+0.32, +1.16] | +1.12 | [+0.63, +1.64] | +3.85 | [+0.77, +7.69] |
| 1.0 | -0.51 | [-0.83, -0.16] | -0.35 | [-0.60, -0.07] | +0.77 | [-3.85, +5.38] |
| 10.0 | -0.88 | [-1.30, -0.46] | -0.91 | [-1.40, -0.40] | -1.54 | [-5.38, +3.08] |
| 100.0 | -0.24 | [-0.54, +0.04] | -0.05 | [-0.62, +0.62] | +2.31 | [-3.85, +9.23] |

**Table 2.** Cross-dataset invariance classification (maximum disagreements per seed across five seeds). Five models are bit-identical across both datasets; four are divergent across both datasets with the same underlying mechanism; two (GaussianNB, SVM_Linear) are empirically near-invariant, differing by at most 1 sample on Kaggle and 0 on OASIS. The structural partition is five structurally invariant and two empirically near-invariant versus four divergent across both datasets.

| Model | Kaggle max disagreements | OASIS max disagreements | Invariance class | Cross-dataset |
| --- | --- | --- | --- | --- |
| AdaBoost | 0 | 0 | Bit-identical | Match |
| HistGradientBoosting | 0 | 0 | Bit-identical | Match |
| KNN | 0 | 0 | Bit-identical | Match |
| SVM Polynomial | 0 | 0 | Bit-identical | Match |
| SVM RBF | 0 | 0 | Bit-identical | Match |
| GaussianNB | 1 | 0 | Near-invariant (empirical) | Bit-id on OASIS |
| SVM Linear | 1 | 0 | Near-invariant (empirical) | Bit-id on OASIS |
| DecisionTree | 215 | 208 | Divergent (RNG-coupled) | Match |
| ExtraTrees | 85 | 81 | Divergent (RNG-coupled) | Match |
| RandomForest | 144 | 111 | Divergent (RNG-coupled) | Match |
| LogisticRegression | 72 | 77 | Divergent (Tikhonov) | Match |

**Table 3.** Logistic Regression color-minus-grayscale effects on both datasets (mean across n=5 seeds per dataset; 95% CIs are paired bootstrap intervals). The direction of effect is preserved across datasets, consistent with the Tikhonov mechanism. The magnitude is smaller on OASIS, consistent with the extent to which L2 regularization affects classification on this data.

| Metric | Kaggle $\Delta$ | Kaggle 95% CI | OASIS $\Delta$ | OASIS 95% CI | Replication |
| --- | --- | --- | --- | --- | --- |
| $\Delta$ Accuracy | +3.55pp | [+3.16, +4.03] | +2.52pp | [+1.65, +3.39] | Match (smaller on OASIS) |
| $\Delta$ Macro F1 | +7.44pp | [+4.76, +10.89] | +5.30pp | [+2.31, +8.32] | Match (smaller on OASIS) |
| $\Delta$ Min-class recall | +26.2pp | [+13.85, +44.62] | +10.77pp | [-3.08, +24.61] | Match (direction; magnitude smaller) |

**Table 4.**
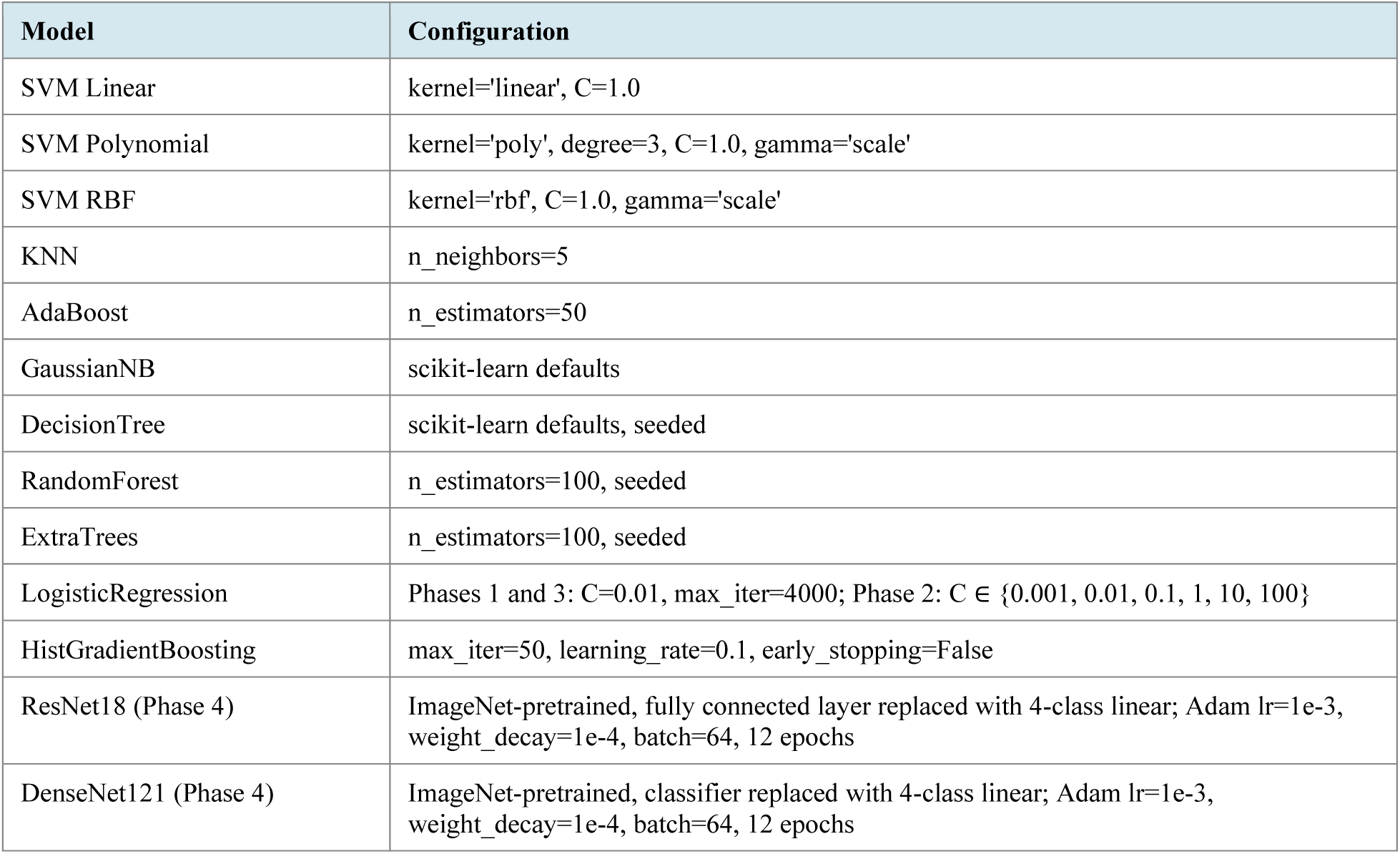
Hyperparameter configuration for all classifiers across all four phases. Classical implementations are from scikit-learn version 1.3.2. CNN implementations are from torchvision (PyTorch 2.1) with the standard ImageNet-pretrained weights.

| Model | Configuration |
| --- | --- |
| SVM Linear | kernel='linear', C=1.0 |
| SVM Polynomial | kernel='poly', degree=3, C=1.0, gamma='scale' |
| SVM RBF | kernel='rbf', C=1.0, gamma='scale' |
| KNN | n_neighbors=5 |
| AdaBoost | n_estimators=50 |
| GaussianNB | scikit-learn defaults |
| DecisionTree | scikit-learn defaults, seeded |
| RandomForest | n_estimators=100, seeded |
| ExtraTrees | n_estimators=100, seeded |
| LogisticRegression | Phases 1 and 3: C=0.01, max_iter=4000; Phase 2: C $\in$ {0.001, 0.01, 0.1, 1, 10, 100} |
| HistGradientBoosting | max_iter=50, learning_rate=0.1, early_stopping=False |
| ResNet18 (Phase 4) | ImageNet-pretrained, fully connected layer replaced with 4-class linear; Adam lr=1e-3, weight_decay=1e-4, batch=64, 12 epochs |
| DenseNet121 (Phase 4) | ImageNet-pretrained, classifier replaced with 4-class linear; Adam lr=1e-3, weight_decay=1e-4, batch=64, 12 epochs |

Figure 7 presents a forest plot summary of color-minus-grayscale effect sizes across all 13 classifiers tested on the Kaggle dataset, and provides a single visual synthesis of the framework’s main empirical claims.

**Figure 7.**
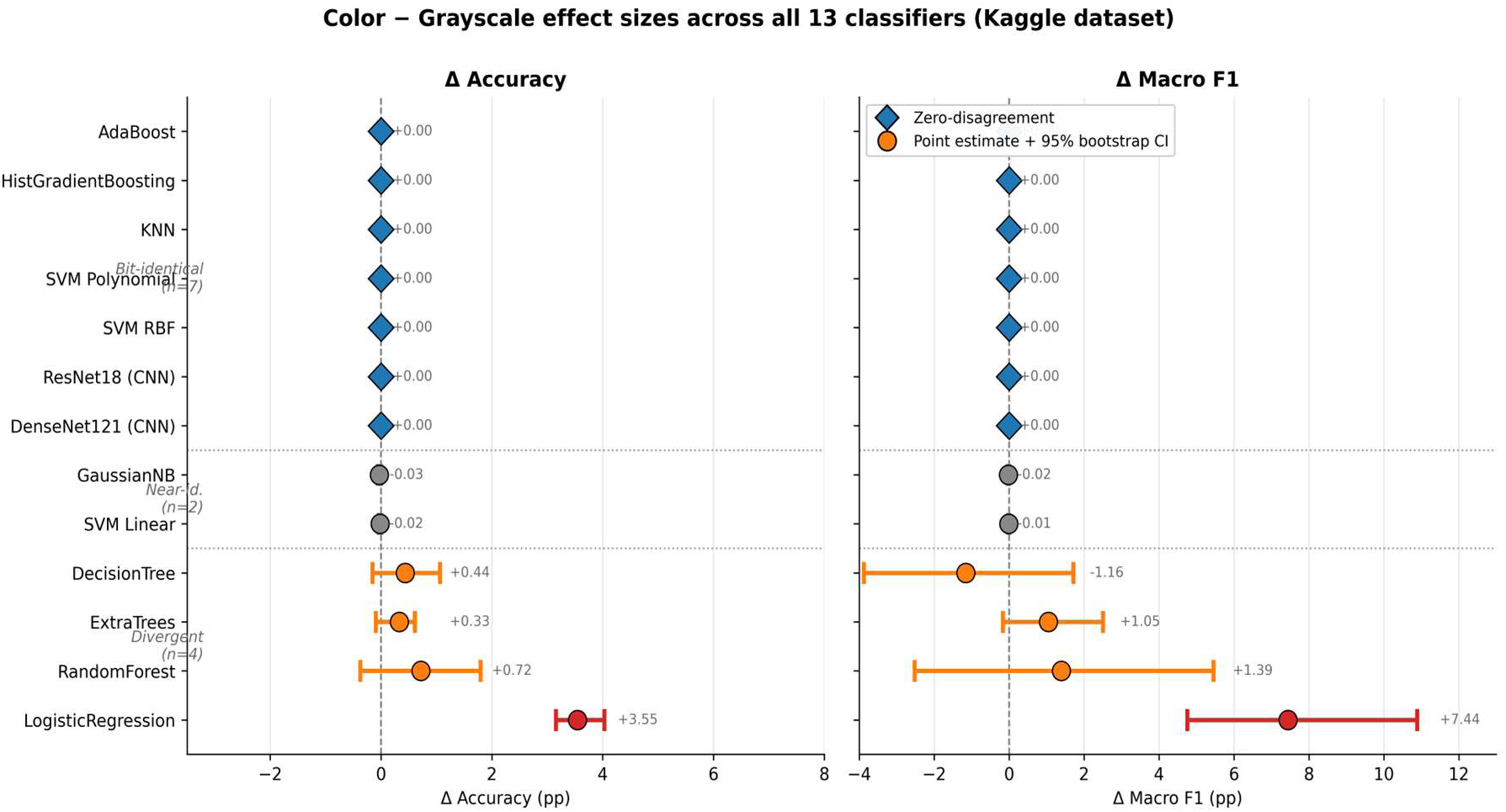
Forest plot of color-minus-grayscale effect sizes for accuracy (left panel) and macro F1 (right panel) across all 13 classifiers on the Kaggle dataset. Diamond markers denote zero-disagreement classifiers (bit-identical group, n=7), where the confidence interval has zero width because predictions match byte-for-byte. Gray circles denote near-identical classifiers (n=2), with effects below 0.05 percentage points reflecting dataset-dependent near-invariance (Section 5). Orange circles denote RNG-coupled divergent classifiers (n=3), whose 95% bootstrap confidence intervals on the difference cross zero, consistent with non-systematic per-seed variation. The red circle denotes LogisticRegression, the only classifier whose 95% CI excludes zero at C=0.01, consistent with the Tikhonov mechanism. The seven bit-identical classifiers include both convolutional networks (ResNet18, DenseNet121), extending the structural-invariance result to deep learning under the standard transfer-learning setup.

## 5. Discussion

The empirical partition presented here uses the mathematical structure of a classifier’s training procedure to predict whether channel triplication of bit-identical RGB data will affect its predictions. Across two datasets, five classifiers were structurally invariant, and four were structurally divergent under channel triplication, with two further classifiers (GaussianNB, SVM_Linear) empirically near-invariant. The five structurally invariant classifiers share a common property: their training procedures and prediction functions depend on quantities that are invariant to the channel triplication operation. KNN’s prediction depends on relative pairwise distances; SVM with polynomial and RBF kernels produces bit-identical predictions specifically because both were run with gamma=’scale’. Triplication triples the pairwise squared distance ‖x−x′‖^2^ (RBF) and the dot product ⟨x,x′⟩ (polynomial), while gamma=’scale’ = 1/(n_features · Var(X)) falls by exactly one-third: the feature count triples, and Var(X) is unchanged because identical copies do not alter the distribution of feature values. The two factors cancel exactly, so every kernel entry is preserved and predictions are bit-identical. This invariance is a property of the scale-gamma default; with gamma fixed to a constant, the kernel arguments would triple, the entries would change, and bit-identity would not hold. AdaBoost with shallow stumps selects single-feature splits whose thresholds are determined by within-feature rankings unaffected by triplication; HistGradientBoosting uses histogram-binned features and tree splits that, when the random state is fixed and no internal feature subsampling enabled, are identical regardless of how many copies of each feature exist. GaussianNB and SVM_Linear are not structurally invariant in the same sense, and no proof of bit-identity is claimed for them. For GaussianNB, triplication multiplies the summed log-likelihood by three while leaving the log-prior unchanged, so the posterior is not uniformly scaled and the argmax can shift; because the 16,384-term log-likelihood dwarfs the small log-prior, the shift only flips predictions at razor-thin margins, which is why one disagreement appears on Kaggle but none on OASIS. For SVM_Linear, distributing weight across three identical feature copies reduces the squared-norm penalty, so triplication is approximately equivalent to scaling the regularization parameter C by three, the same Tikhonov coupling identified for Logistic Regression. SVM_Linear appears near-invariant here only because it was run at C=1, in the hinge-dominated regime where a threefold change in effective C barely moves the solution; a regularization sweep analogous to Phase 2 may reveal a similar monotonic gap and is recommended as a confirmatory analysis.

The four divergent classifiers diverge through two distinct mechanisms. DecisionTree, RandomForest, and ExtraTrees diverge through random-number-generator coupling to the feature count, by two related routes. RandomForest and ExtraTrees draw a random subset of candidate features at each split (max_features=’sqrt’), so the subset drawn from a 49,152-element index space differs from the equivalent draw on a 16,384-element space even with the random state fixed. DecisionTree at scikit-learn defaults (max_features=None) does not subsample features; every split evaluates all features, so no information is lost. Choosing between two identical copies of a feature yields the same partition and cannot by itself change a prediction. Its divergence is instead a tie-breaking effect: the splitter visits features in an RNG-shuffled order, and that order is consumed differently at 49,152 versus 16,384 features, so ties among genuinely distinct, equal-improvement splits are resolved differently. Its divergence is therefore a permutation/tie-breaking effect rather than subsampling. This mechanism produces non-systematic per-seed differences with confidence intervals on the mean effect spanning zero. LogisticRegression diverges via Tikhonov coupling, where the L2 regularization penalty scales with the feature count. With feature triplication, the optimum can distribute weight across the three identical copies, with each copy carrying one-third of the magnitude and contributing one-ninth of the penalty. The effective regularization is therefore relaxed by a factor proportional to the feature multiplication. The Phase 2 regularization sweep, a controlled interventional ablation, demonstrates this mechanism within the tested implementation: as the regularization parameter C increases (regularization weakens), the color-minus-grayscale gap shrinks from +12 percentage points on macro F1 at C=0.001 to within 0.05 percentage points of zero at C=100. The mechanism is not a property of the data; it is a property of the optimization objective.

The minority-class effect is particularly notable. The Moderate Demented class is 1% of the Kaggle dataset, with 13 test samples per seed. At C=0.01, strong L2 regularization on grayscale shrinks minority-class weights toward zero, producing a mean recall of 0.65 with high seed-to-seed variance. Under color, the effectively relaxed regularization allows the model to maintain larger minority-class weights, raising mean recall to 0.91 while reducing variance. The 26.2 percentage point recall improvement on the Kaggle dataset is the largest practical effect of channel triplication observed in this study. Phase 2 confirms that this effect is entirely a regularization artifact: at C=100, the same color-versus-grayscale comparison produces no minority-class advantage. Phase 3 confirms that the direction of the effect is robust to dataset choice: on OASIS, the same Logistic Regression at C=0.01 produces a 10.77 percentage-point improvement in minority-class recall for color over grayscale.

The cross-dataset replication also resolves the status of the near-identical group from Phase 1. On the Kaggle dataset, GaussianNB and SVM_Linear each produced a single disagreement in their predictions across one or two seeds. The Phase 3 OASIS results showed no disagreements for either classifier across all five seeds. Neither classifier is provably bit-identical under channel triplication (Section 5): for GaussianNB, the log-prior is left unscaled while the log-likelihood triples; for SVM_Linear, the L2 penalty couples to the feature count, as in Logistic Regression. Both effects are small in these datasets, so the Kaggle disagreement amounts to one sample near a class-margin boundary, whereas OASIS shows none. The structural partition is therefore five invariant versus four divergent, with GaussianNB and SVM_Linear forming a separate, empirically near-invariant pair rather than members of a proven bit-identical group.

The findings have practical consequences for reproducibility audits and the design of benchmarks in medical imaging machine learning. Logistic Regression was tested directly, and the Tikhonov coupling was observed; similar behavior is expected in other L2-regularized linear models on bit-identical RGB data, since the underlying mechanism (regularization penalty scaling with feature count) is a property of the L2 objective rather than of Logistic Regression specifically. Reproducibility audits should report whether channel triplication was applied to classical-ML pipelines using such models, and whether regularization-aware ablations were performed. For deep convolutional networks in the standard transfer-learning setup (ImageNet-pretrained model with three-channel input fine-tuned on triplicated grayscale data), the Phase 4 result establishes that channel triplication is invariant at the prediction level and identical in computational cost: the choice between loading images as grayscale-then-expanded or as native triplicated RGB is arbitrary at the prediction level and has no compute consequences. The classical ML compute waste finding (channel triplication multiplies training time by 2.3 to 4.0 with no benefit for the bit-identical group) does not apply to CNNs because both code paths produce the same input tensor shape. The overall classifier-family picture is therefore: for five structurally invariant classical models, two further empirically near-invariant models, and both convolutional networks tested, channel triplication is a no-op or near-no-op at the prediction level; for four of the eleven classical models, it produces measurable changes through identified mechanisms; for the four divergent classical models specifically, color-condition compute cost is 2.3 to 4.0 times higher than grayscale, with Logistic Regression’s gain being a regularization artifact rather than an information gain.

## 6. Limitations

### 6.1 Two Datasets, Both Alzheimer’s MRI

The empirical findings are based on two Alzheimer’s MRI datasets (Kaggle and OASIS). The theoretical mechanisms identified (bit-identity via invariant decision functions, stochastic divergence via feature subsampling, Tikhonov coupling via L2 regularization scaling) apply to any classifier evaluated on bit-identical triplicated channels, regardless of the underlying modality. However, empirical demonstration across additional medical imaging modalities (e.g., chest X-ray, histopathology, dermatology) would further strengthen the external validity claim.

### 6.2 CNN Configuration Tested

Phase 4 tested ResNet18 and DenseNet121 in the standard transfer-learning configuration: ImageNet-pretrained, three-channel input convolution, fine-tuned end-to-end on Alzheimer’s MRI data with channel-triplicated grayscale input. The bit-identical finding follows from the input tensor identity under this configuration. A different configuration could produce different results. Custom CNNs with first-layer convolutions modified to accept single-channel input would not be tested in this experiment, because the grayscale-versus-color comparison would lack a well-defined meaning in that setup (the architectures themselves would differ). Vision transformers that patch-embed inputs differently would also require separate evaluation. The Phase 4 result establishes the bit-identical group for the dominant deep-learning paradigm in medical imaging, not for all possible deep-learning configurations.

### 6.3 Image Resolution

All experiments used 128 × 128 pixel images. Higher resolutions increase the feature count proportionally and would not change the qualitative findings, but the magnitude of the Tikhonov effect on linear models may scale with resolution.

### 6.4 Minority Class Sample Size

The Moderate Demented class contains 64 images in each dataset (Kaggle and the OASIS subsample), yielding 13 test samples per seed. The 26.2 percentage-point recall improvement observed on the Kaggle dataset is based on a small number of test samples, and the confidence interval is correspondingly wide. The relative improvement and direction are consistent across seeds and replicates on OASIS, supporting the qualitative finding.

## 7. Future Work

Several directions could extend this work. First, replication across additional medical imaging modalities would test the framework’s generality beyond MRI, with chest X-rays, histopathology images stored as RGB, and dermatology datasets as natural targets. Second, evaluating CNN configurations outside the standard transfer-learning setup, such as custom single-channel input convolutions, vision transformers with patch embedding, and hybrid architectures, would extend the deep-learning analysis to settings where the input-tensor-identity argument no longer holds. Third, formal proofs of the bit-identity property for each relevant classifier family would strengthen the theoretical foundation. Fourth, audit tools that automatically detect channel triplication in published medical imaging pipelines and flag the affected classical-ML results would have direct practical value for reproducibility research.

## 8. Conclusion

Channel triplication is a common preprocessing step in medical imaging machine learning pipelines, and it is applied largely without justification or audit. This research tested its consequences directly across four coordinated experiments. Phase 1 partitioned 11 classical machine learning models into three empirical groups on the Kaggle Alzheimer MRI Image Dataset: bit-identical (five models), near-identical (two models), and divergent (four models). Phase 2 provided strong evidence for the Tikhonov mechanism through a six-point regularization sweep on Logistic Regression, in which the color-minus-grayscale macro F1 gap shrank monotonically from +12 percentage points under strong regularization to within 0.05 points of zero under weak regularization, consistent with the proposed coupling between L2 regularization and feature count. Phase 3 replicated the Phase 1 partition on the OASIS Alzheimer’s Detection dataset, with one refinement: the two Kaggle-near-identical models produced zero disagreements on OASIS, showing that the near-identical pair is empirically near-invariant and dataset-dependent rather than structurally identical. The Logistic Regression Tikhonov effect was also replicated on OASIS, in the same direction but with a smaller magnitude, again consistent with the mechanism. Phase 4 extended the analysis to ResNet18 and DenseNet121 across both datasets: all 20 paired conditions produced byte-identical predictions, placing ImageNet-pretrained convolutional networks with channel-triplicated input in the bit-identical group.

The practical picture that emerges is straightforward. For five of the eleven classical models, channel triplication is byte-identical at the prediction level; two further models are empirically near-identical; and both convolutional networks tested are byte-identical. For the four divergent classical models, it alters predictions, producing classifier-dependent effects through the mechanisms identified above: random-number-generator coupling to the feature count in the tree ensembles, and L2 regularization coupling in Logistic Regression. On cost, triplication multiplies training time by 2.3 to 4.0 times for the classical models, with no benefit, while leaving training time unchanged for the convolutional networks, where both code paths produce the same input tensor. Researchers who report accuracy improvements from color input on grayscale-equivalent medical imaging data should verify that the gain is not a Tikhonov artifact from a regularized classical model.

## Declarations

Competing interests: The authors declare no competing interests. Funding: This research received no external funding. Corresponding author: Sanjay Singhvi, CompuTerra Inc. Use of AI tools: The authors used an AI assistant (Anthropic Claude) to support manuscript editing and revision; the authors are responsible for all content, including the study design, experiments, analyses, and conclusions, and reviewed and verified the manuscript in full. Ethics and scope: This study uses publicly available image datasets and does not involve new human-subject data collection. The analysis audits preprocessing effects in machine-learning pipelines and should not be interpreted as clinical diagnostic validation. Author contributions: R.S. and S.S. designed the study, implemented the experiments, analyzed the results, and wrote the manuscript.

## Data and Code Availability

The Kaggle Alzheimer MRI Image Dataset is publicly available at https://www.kaggle.com/datasets/sachinkumar413/alzheimer-mri-dataset. The OASIS Alzheimer’s Detection dataset is publicly available at https://www.kaggle.com/datasets/ninadaithal/imagesoasis. The OASIS subsample manifest (6,400 file paths selected from the larger dataset using seed=0), the 110-cell Phase 1 experimental data, the 120-cell Phase 2 regularization sweep data, the 110-cell Phase 3 replication data, the 40-cell Phase 4 CNN data, per-cell prediction arrays, channel identity verification logs, and figure-generation scripts are available from the corresponding author on request. The full reproducible pipeline is implemented as four Python notebooks (Phase 1: CPU; Phase 2: CPU; Phase 3: CPU; Phase 4: GPU T4 x1) suitable for headless execution on Kaggle, with automatic resume from any prior partial run and complete reproducibility via fixed seeds throughout.

